# Doderlin: Isolation and Characterization of a Broad-Spectrum Antimicrobial Peptide from *Lactobacillus acidophilus*

**DOI:** 10.1101/2022.01.19.476933

**Authors:** Bruna S. da Silva, Andrea Díaz-Roa, Erica S. Yamane, Mirian A. F. Hayashi, Pedro Ismael da Silva Junior

## Abstract

*Lactobacillus acidophilus* are Gram-positive bacteria distributed in diverse environments, and as being a component of the normal microbiota of gastrointestinal and urogenital tract, being relevant to humans. Classified as lactic acid bacteria, due to the pro-duction of lactic acid, *Lactobacillus* can also produce antimicrobial peptides (AMPs), which is a compound synthesized by all forms of life aiming for protecting themselves from threats and to increase their competitivity to survive in a specific environment. AMPs are molecules capable of inhibiting the growth of microorganisms and, due to the indiscriminate use of conventional antibiotics and the emergence of multi-resistant bacteria, they have become an alternative, not only for treating multi-resistant infections, but also for probiotic product confection and food conservation. Considering the rampant rise of resistance, the present study aimed to isolate and characterize antimicrobial peptides from *Lactobacillus acidophilus* extracts. Samples were obtained from *Lactobacillus* acid extract supernatant which was pre-fractionated on disposable cartridges, followed by a high-performance liquid chromatography (HPLC). The collected fractions were evaluated in a liquid growth inhibition assay where eight fractions antimicrobial activity were obtained. One of them was selected for further characterization by mass spectrometry (MS), due to its antimicrobial activity against *Candida albicans* and conclusive results in mass spectrometry analysis. This molecule was identified as a peptide having a molecular mass of 1788.01 Da, peptide sequence NEPTHLLKAFSKAGFQ, and named Doderlin. Interestingly, antimicrobial molecules isolated from *L. acidophilus* have already been described previously, but few reports in the literature describe an AMP effective against *C. albicans* as reported here. The results obtained suggest that this newly discovered molecule have a biological property with potential to be applied in pharmaceutical and food companies in the fight against contamination and/or for treating infections caused by microorganisms.

**IMPORTANCE:** Doderlin, this newly discovered molecule have a biological property with potential to be applied in pharmaceutical and food companies in the fight against contamination and/or for treating infections caused by microorganisms.

## Introduction

The lactic acid bacteria are a very diverse group of Gram-positive fermenting human-associated bacteria as members of their normal microbiota, some of them also known as probiotics (1). As typical characteristics this bacteria is a catalase negative, non-spore-forming anaerobic, cocci or bacillus format, lacking locomotion structures and cytochromes (2, 3). Among the benefits of using probiotics are the aid in preventing diarrhea, infectious diseases and stomach ulcers, decreased lactose intolerance and allergies, stimulation of intestinal and systemic immunity, antimicrobial activity, inhibitory action against some cancers and control of cholesterol levels (4–6).

The beneficial effects of *Lactobacillus* come from the production of a large number of active metabolites which assist them in rivalry with other microorganisms that occupy the same ecological niche. Among these metabolites are: organic acids (lactic acid, acetic acid and other short chain), hydrogen peroxide, carbon dioxide, low molecular weight antimicrobial substances, bacteriocins and bacteriocin-like substances (7). These metabolites, especially hydrogen peroxide and lactic acid, from *Lactobacillus* are also routinely employed in many industrial processes including the production of fermented foods such as vegetables, meats and dairy products (milk, yogurt and cheese) (8, 9).

Among the species belonging to the group of *Lactobacillus* stand out the *Lactobacillus acidophilus*. These bacteria are present in the human oral, gastrointestinal and urogenital mucosa, where they adhere to the epithelium and alter their environment in such a way that colonization by other bacteria or fungi through competition for resources, predation by bacteriophage and production of inhibitory metabolites is difficult (10–12).

Also known as Döderlein’s bacilli, *L. acidophilus* was first described by Professor Albert Döderlein in 1892. In his pioneering studies on bacteria present in vaginal secretions, Döderlein characterized such organisms as Gram-positive, fermenter, non-spore-forming and unflagged bacteria (10, 13, 14). He also noted that the bactericidal activity of vaginal secretions was associated with lactic acid production due to the fermentation process performed by the bacilli (15). *L. acidophilus* ais the most abundant bacteria in these environments, and a member of the normal gut and vagina microbiota (16, 17). Its role in the human intestinal microbiota is to assist the level of harmful bacteria and fungi lowering and to produce lactase, which is an important enzyme in milk digestion (18). The presence of lactic acid, produced by both *L. acidophilus* and the vaginal mucosa, lowers the local pH to about 4.0 and 4.5, creating an environment that limits the growth of other bacteria (14). *L. acidophilus*, as well as all *Lactobacillus*, basically present two ways of interference against pathogens: (1) adherence to the epithelium forming a barrier that prevents colonization, leading to competition for receptors present in epithelial cells, or (2) the production of antimicrobial compounds, such as acids hydrogen peroxide, bacteriocins and antimicrobial peptides (AMPs) (10, 19, 20).

AMPs compose a class of compounds highly heterogeneous that are generally defined as small (cationic, ranging from 5 to 100 amino acid residues) molecules with high proportion of hydrophobic residues and high-water solubility, which are properties that allow the necessary interactions of AMPs to activate their mechanisms of action (21, 22).

Native AMPs are one of the most important defense systems in either prokaryotic or eukaryotic organisms, showing a highly diverse composition, and a dual anti-inflammatory and antimicrobial effect has been described for the most known AMPs. Thousands of native and chemically synthesized AMPs were discovered and reported (23, 24). In prokaryotes, which have a simpler structure compared to eukaryotes and with no immune system, AMPs confer an ecological advantage in competition for resources necessary for survival, especially with a strong activity against organisms that occupy a similar niche or habitat (25).

The typical mechanism of action of an AMP usually involves the electrostatic interaction with the plasma membrane, followed by the formation of pores that lead to intracellular content leakage and, consequently, to the cell death (26–28). However, some peptides can interact to intracellular targets, like nucleic acids, as suggest in *in vitro* studies with Sarconesin II (29) and Rondonin (30).

The indiscriminate use of antibiotics and the emergence of multidrug-resistant bacteria has rekindled the interest in AMPs, which was overshadowed by the era of antibiotics that began in 1928 with Alexander Fleming’s discovery of penicillin (31). Antibiotics were a great discovery of the twentieth century, but they are losing their effectiveness increasingly due to the evolution of microbial resistance mechanisms (32). This topic is so worrisome that it made the United Nations include it on the list of greatest health threats in 2019.

In this context, the present work is focused on bioprospection and functional and structural characterization of a novel short peptide. Natural sources, including bacteria, are excellent samples for this purpose. Despite its moderate and broad-spectrum antimicrobial activity, this study presents a unique sequence that can be modified, and explored in future structure-function studies. Additionally, the selectivity evidenced by the absence of hemolytic activity at high concentrations is a determining factor for translation to in vivo studies.

## RESULTS

### Identification and isolation of the active fraction/compound from the Lactobacillus acidophilus extracts

Several fractions were eluted from *Lactobacillus acidophilus* extracts which were all submitted to the liquid growth inhibition assay against Gram-positive, Gram-negative, and yeasts, aiming for the identification of the fractions with antimicrobial activity. Only the samples eluted at 5 and 40% ACN concentration presented antimicrobial activity. The fractions eluted at 80% ACN did not show any antimicrobial activity. A total of eight fractions inhibited the growth of the tested microorganisms. Four fractions were eluted at 5% ACN (acetonitrile), numbered 3, 12, 14, and 15 (Figure 1A), while other four were eluted at 40% ACN: 23, 25, 29, and 32 (Figure 2A). The corresponding antimicrobial fractions and the microorganisms inhibited by them are depicted summarized in Table 1.

**Figure 1.**
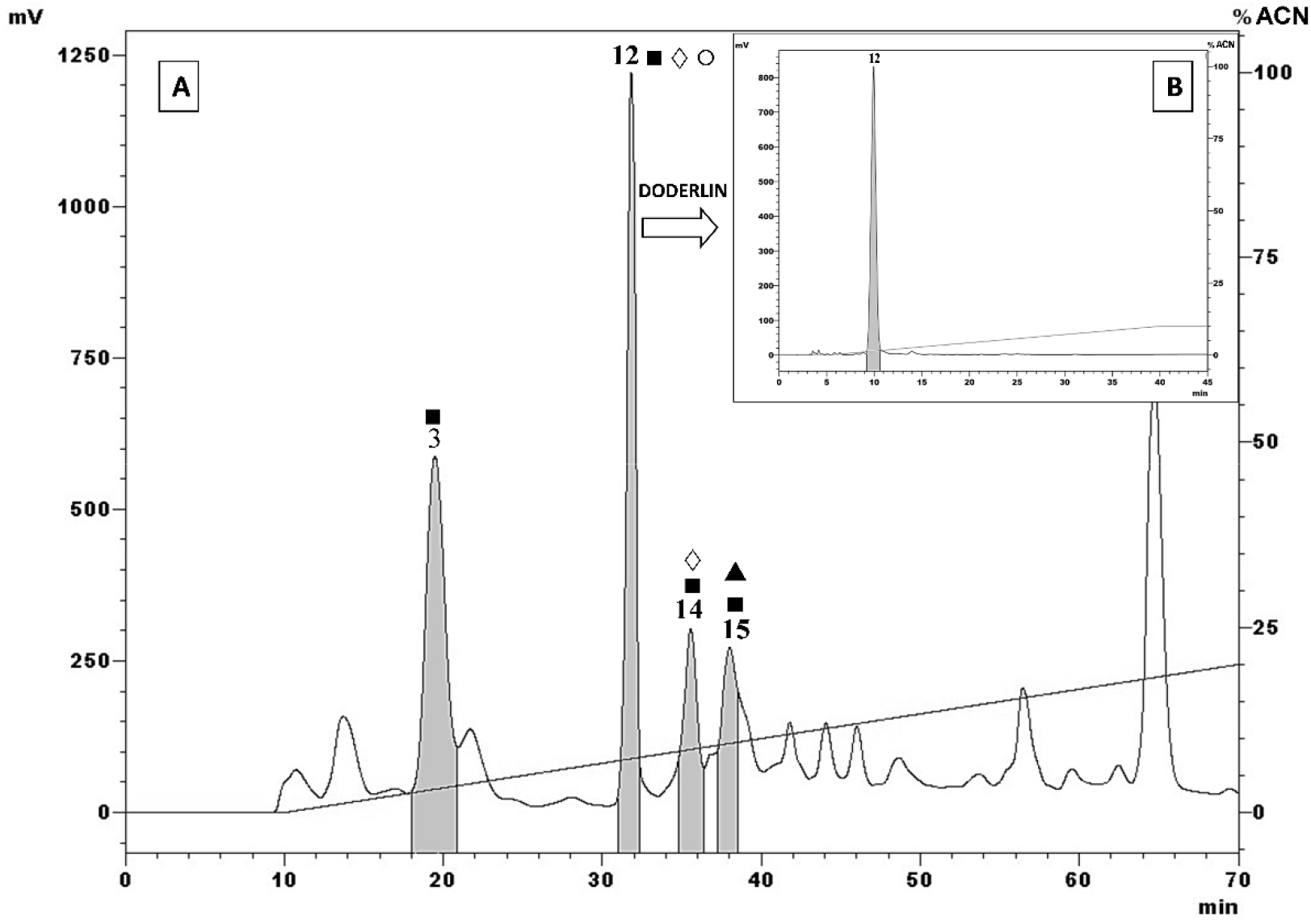
Reverse-phase high-performance liquid chromatography (RP-HPLC) profile *of Lactobacillus acidophilus* extract eluted at 5% ACN on the Sep-Pak^®^ pre-fractionation step. (**A**). Fractions eluted on a Jupiter^®^ C18 semi-preparative column (10 × 250 mm, 10 μm; 300 Å), with a linear gradient 0%-20% ACN in acidified water for 60 min, at a flow rate of 1.5 mL/min, monitored at 225 absorbance. The fractions were manually collected and submitted to the antimicrobial liquid growth inhibition assay. Peaks highlighted indicate antimicrobial activities against the tested microorganisms: *Microccocus luteus* A270 (∎), *Escherichia coli* SBS 363 (▲), *Pseudomonas aeruginosa ATCC27853* (@), and *Candida albicans* MDM 8 (○). The numbered peak 12 corresponds to the Doderlin sample eluted at 30.8-32.3 min, and resubmitted to the system using a Jupiter^®^ C18 analytical column (4.6 mm × 250 mm, 10 μm; 300 Å), with a linear gradient 0%-10% ACN in acidified water for 40 min, at a flow rate of 1 mL/min. The absorbance was monitored at 225 nm (**B**).

**Figure 2.**
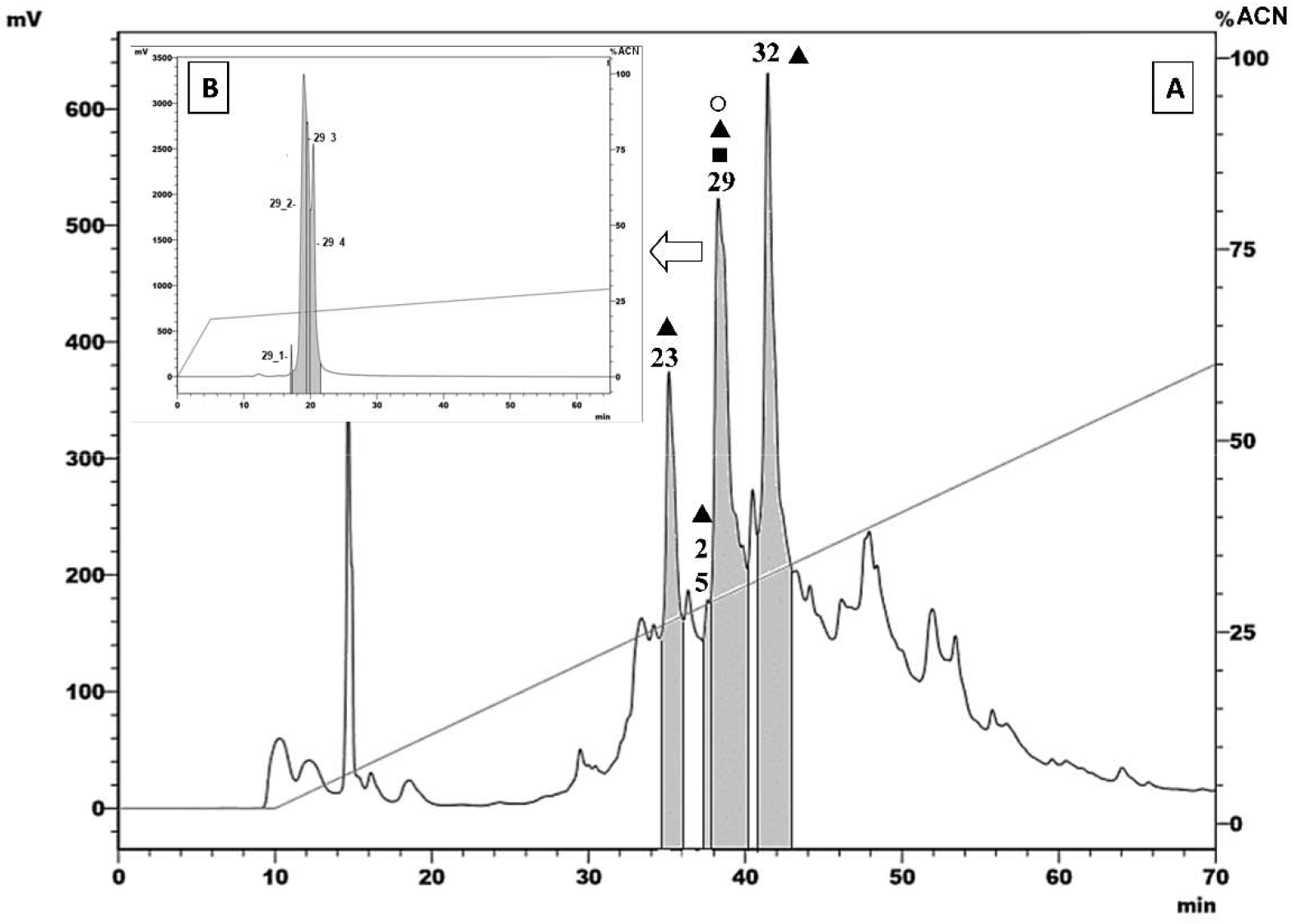
Reverse-phase high-performance liquid chromatography (RP-HPLC) profile *of Lactobacillus acidophilus* extract eluted at 40% ACN on the Sep-Pak^®^ pre-fractionation step. (**A**). Fractions eluted on a Jupiter^®^ C18 semi-preparative column (10 × 250 mm, 10 μm; 300 Å), with a linear gradient 0-60% ACN in acidified water for 60 min, at a flow rate of 1.5 ml/min, monitored at 225 absorbance. The fractions were manually collected and submitted to the antimicrobial liquid growth inhibition assay. Peaks highlighted indicate antimicrobial activities against the tested microorganisms: *Microccocus luteus* A270 (∎), *Escherichia coli* SBS 363 (▲), and *Candida albicans* MDM 8 (○). The numbered peak 29 corresponds to the sample eluted at 37.9-40.3 min, was resubmitted to the system using a Jupiter^®^ C18 analytical column (4.6 × 250 mm, 10 μm; 300 Å), with a linear gradient 19-29% ACN in acidified water for 60 min, at a flow rate of 1 mL/min. The absorbance was monitored at 225 nm (**B**).

**Table 1.**
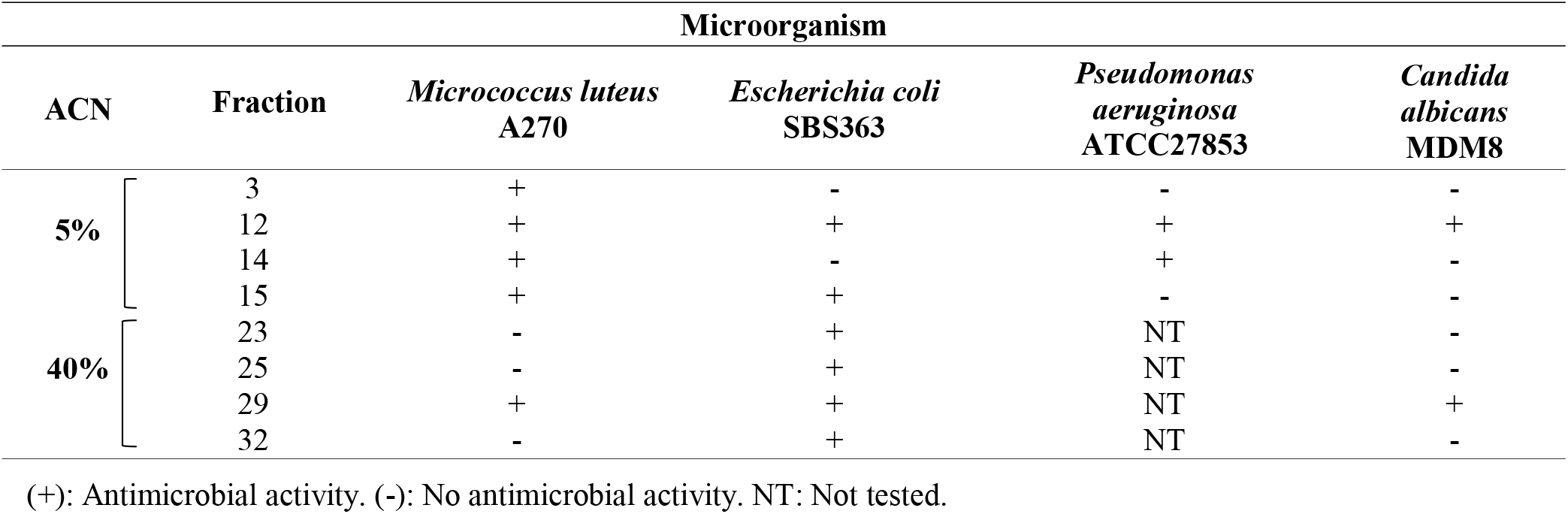
Antimicrobial activity of all detected fractions.

| ACN | Fraction | Microorganism |  |  |  |
| --- | --- | --- | --- | --- | --- |
|  |  | *Micrococcus luteus*<br>A270 | *Escherichia coli*<br>SBS363 | *Pseudomonas aeruginosa*<br>ATCC27853 | *Candida albicans*<br>MDM8 |
| 5% | 3 | + | - | - | - |
|  | 12 | + | + | + | + |
|  | 14 | + | - | + | - |
|  | 15 | + | + | - | - |
| 40% | 23 | - | + | NT | - |
|  | 25 | - | + | NT | - |
|  | 29 | + | + | NT | + |
|  | 32 | - | + | NT | - |

(+): Antimicrobial activity. (-): No antimicrobial activity. NT: Not tested.

Fractions identified as having antimicrobial potential were carried out to a second chromatography step in order to verify the homogeneity of the samples. In this step, the samples were selected based on the antimicrobial activity against the yeast *Candida albicans*, which is an opportunistic fungus of medical relevance affecting the human female genital tract, and the fractions 12 (eluted at 30.8-32.3 min), and 29 (eluted at 37.6 min) were then selected. Fraction 12 (Figure 1B) was apparently homogeneous, while from the fraction 29 were eluted four main peaks (Figure 2B). These samples were submitted again to the liquid growth inhibition assay to ensure that the antimicrobial activity remained even after this new fractionation process, and only fraction 12 was homogeneous and conserved the same antimicrobial activity observed in the first assay (Table 1). Therefore, it was selected for the characterization steps.

### Mass spectrometry (MS) and characterization of the compound with antimicrobial activity

The mass spectrometry (MS) analysis of the fraction 12 highlighted the prominent ion corresponding to a molecular weight (MW) of 664 Da besides other ions, indicating a probable MW above 1 kDa (Figure 3). Due to the presence of these ions, a new step of liquid chromatography was performed using a gel-filtration column to separate the samples by their size.

**Figure 3.**
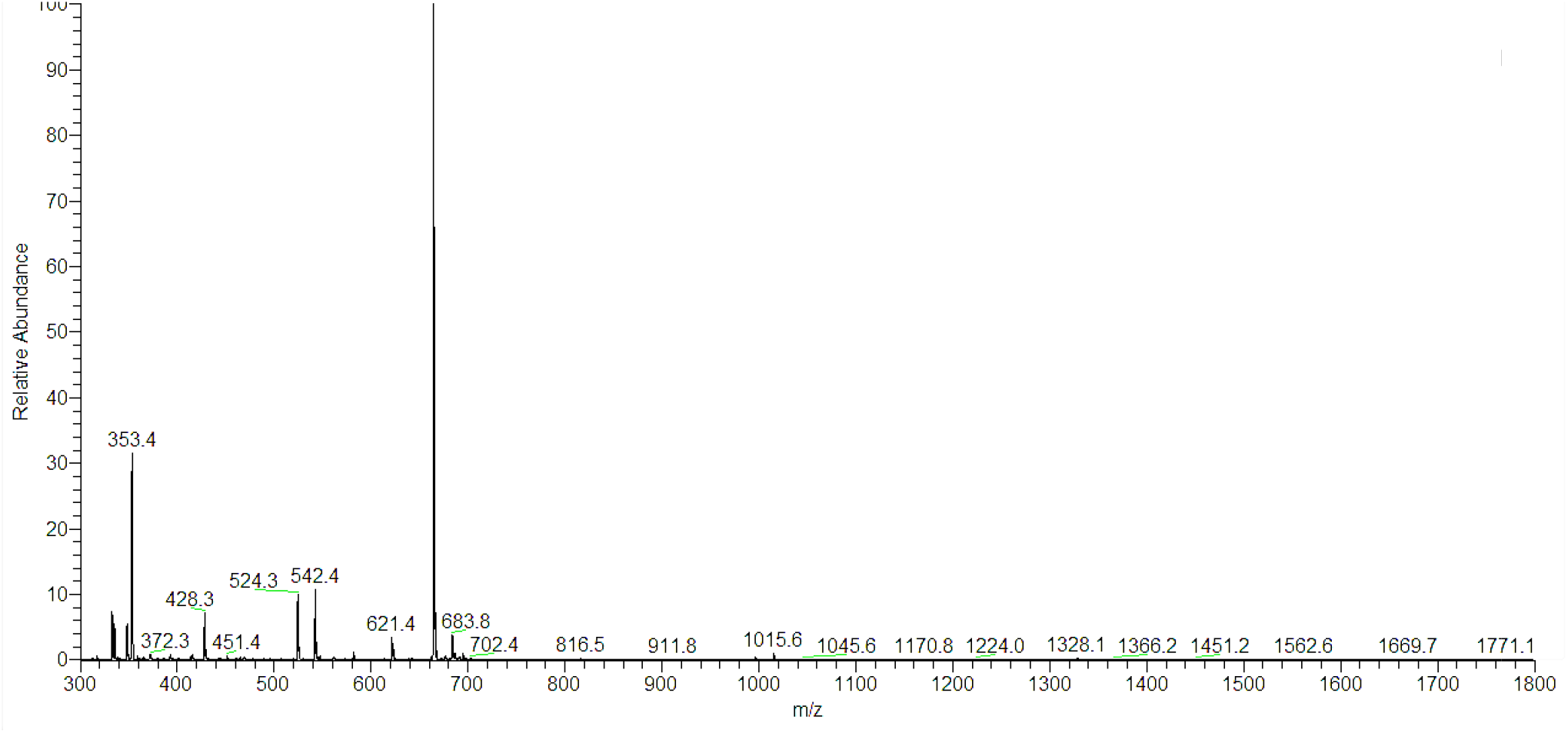
Full MS spectrum of fraction 12 generated by Xcalibur^®^ software. Obtained spectrum evidencing the most prominent ion with 664.4 m/z (mass/charge ratio), and visible ions with higher than 1,000 m/z, indicating that the molecule mass could be more than 1 kDa.

This new chromatographic profile detected two different fractions which were eluted from the fraction 12, numbered 1 and 2 (Figure 4). Both samples were subjected to MS analysis again, allowing to identify one fraction (1) with a molecular weight of 664 Da m/z (already identified in the first MS step, Figure 5B) and the other, fraction (2), corresponding to a molecule with 1,788 Da m/z (Figure 5A). Both samples were retested for the antimicrobial activity, but only the 1,788 Da molecule presented antimicrobial activity similar to that described initially in Table 1.

**Figure 4.**
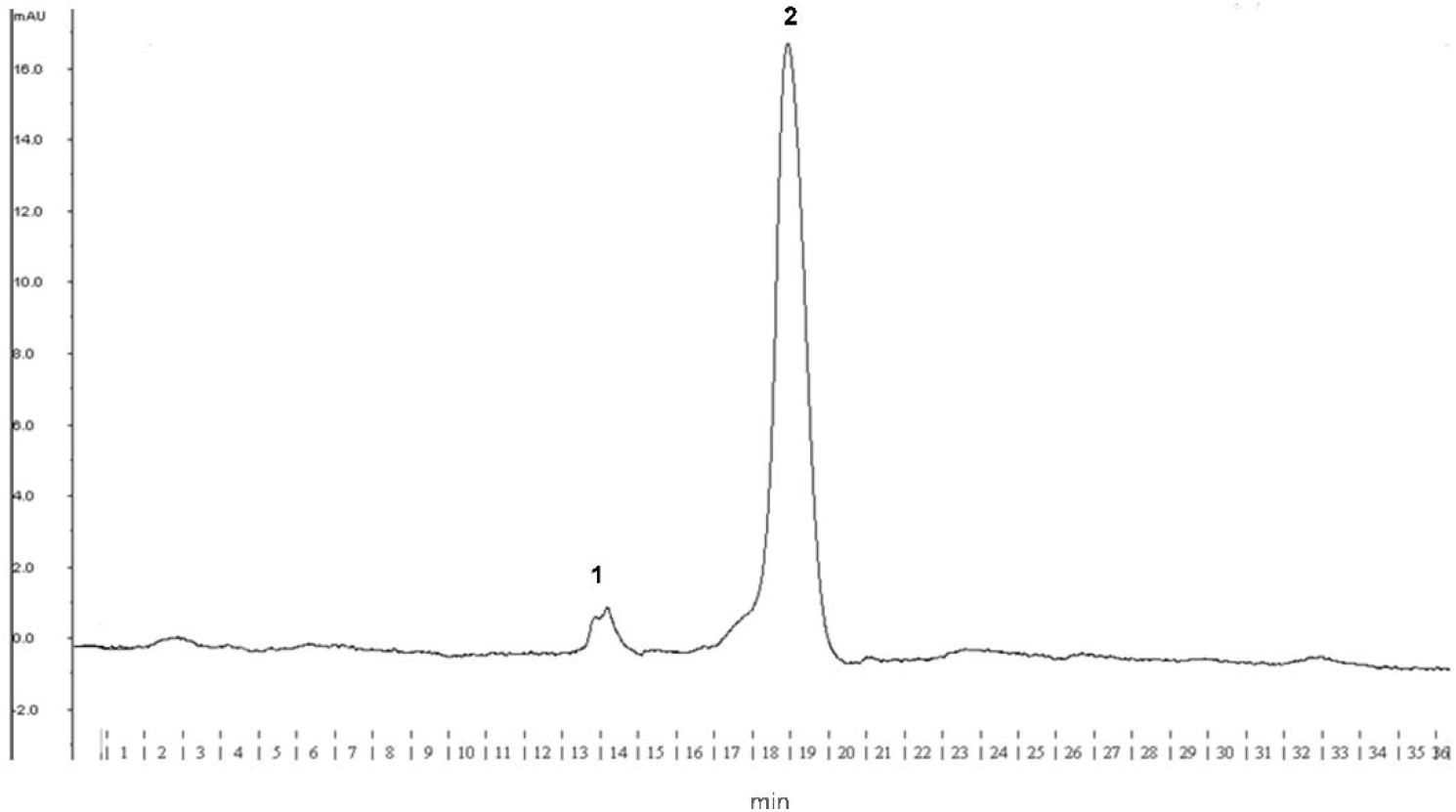
Size-exclusion chromatography (SEC) profile of fraction 12. Peaks eluted after sample submission to SEC system, carried out on a Superdexpeptide HR 10/30 column, under isocratic conditions, using a 50 mM ammonium acetate solution as eluent. The absorbance was monitored at 280 nm. Fractions were collected automatically every minute for 36 min.

**Figure 5.**
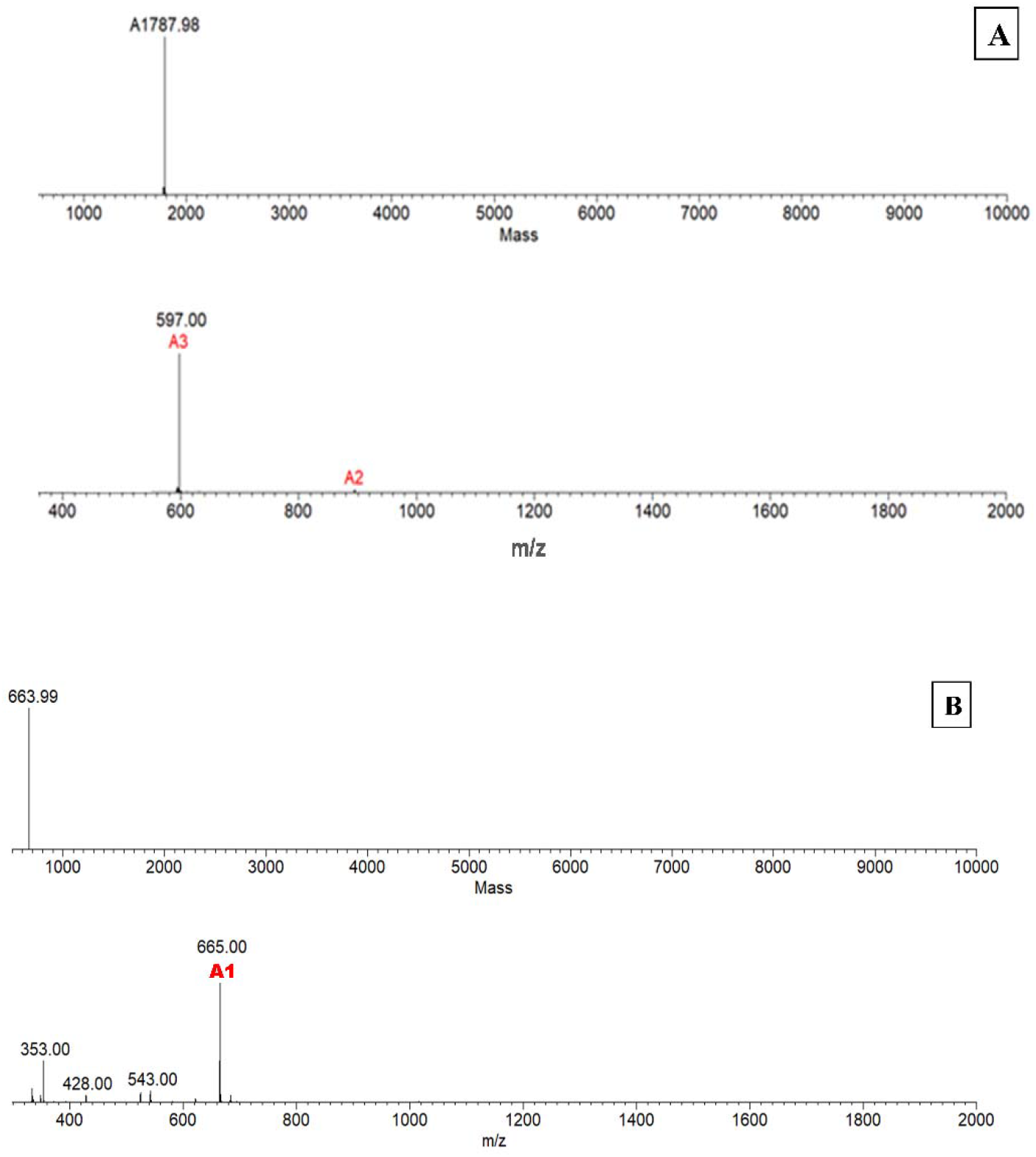
Fraction 12 ion deconvolution and molecular mass generated by Magtran software after fractionation by SEC. Mass spectrometry analyses revealed a molecular mass of approximately 1,788 Da for Doderlin, based on mass/charge (m/z) of its double-charged ion [M + 2H]^2+^, and triple-charged ion [M + 3H]^3+^, where A2 (895 m/z) correspond to double-charged ion, and A3 (597 m/z) to triple-charged ion (**A**); A1 (665 m/z) correspond to single-charged ion [M + H]^+^, revealing the same 664 Da molecule identified before SEC step (**B**).

Analysis using the in-house MASCOT tool allowed identifying the primary sequence NEPTHLLKAFSKAGFQ for this molecule (Figure 6A), which was drawn using the PepDraw tool as shown in (Figure 6B). This sequence was named Doderlin, in honor to the responsible for the discovery of the vaginal bacilli Dr. Albert Döderlein. In addition, as the native purified molecule isolated from the *Lactobacillus acidophilus* extracts has limited quantitative recovery due to the several fractionation steps necessary for the complete purification process, the full-length analog was chemically synthesized and commercially purchased.

**Figure 6.**
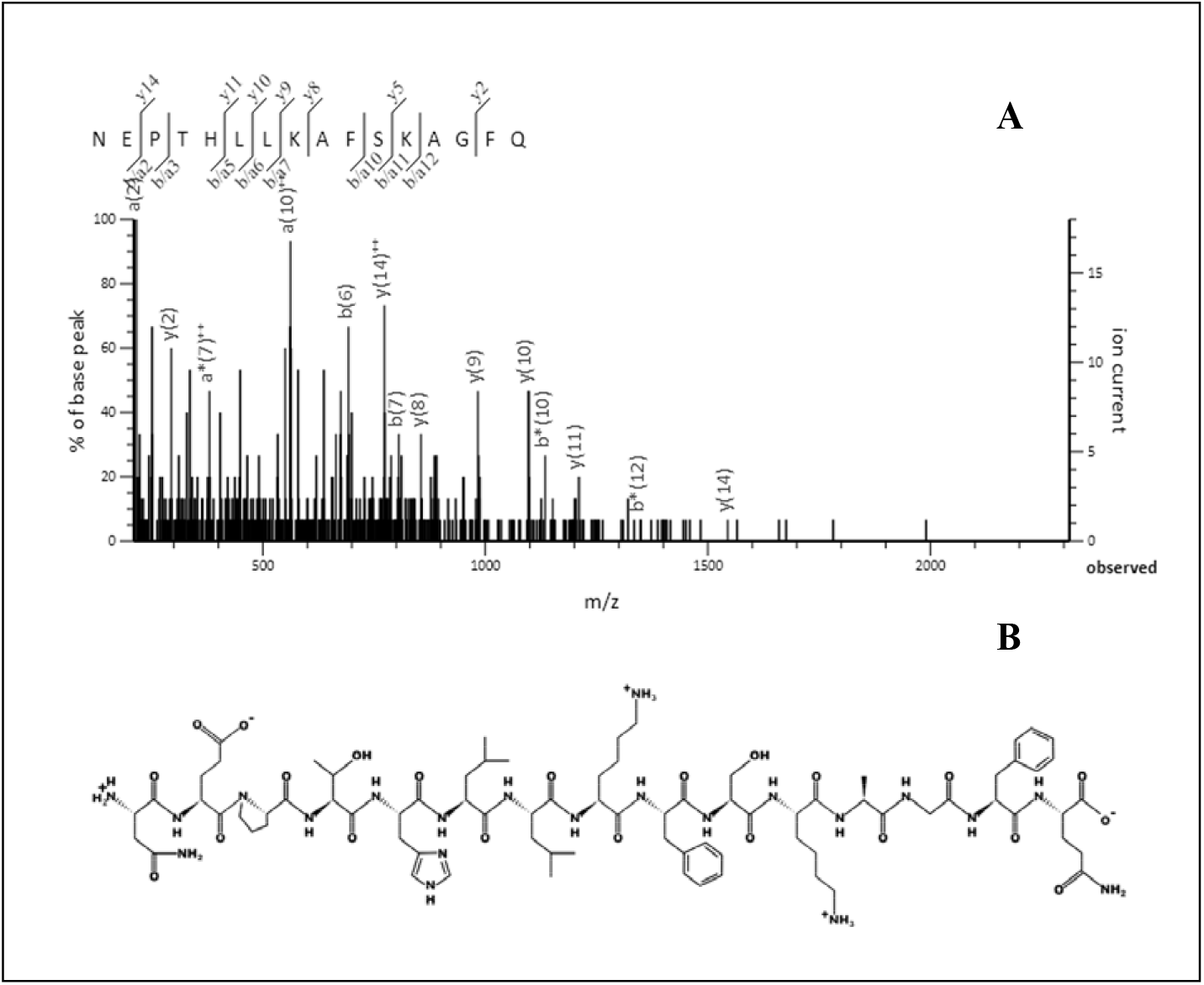
Doderlin peptide sequence generated by Mascot tool. Representative MS/MS spectra generated from CID fragmentation of Doderlin and submitted to Mascot search, revealing the “NEPTHLLKAFSKAGFQ” sequence denoted by deconvolution of “b” and “y” peptide ions (**A**). The primary structure of the Doderlin peptide generated by Pepdraw software (**B**).

### Bioassays

#### Antimicrobial spectrum analysis of chemically synthesized Doderlin

A liquid growth inhibition assay was performed again with the chemically synthesized Doderlin to confirm and extend the knowledge on the antimicrobial spectrum of this peptide.

Among the microorganisms evaluated here, Doderlin was ineffective against most Gram-negative bacteria and filamentous fungi tested here (Table 2). On the other hand, Doderlin showed a wide spectrum of antimicrobial action against Gram-positive bacteria and yeasts (Table 2).

**Table 2.**
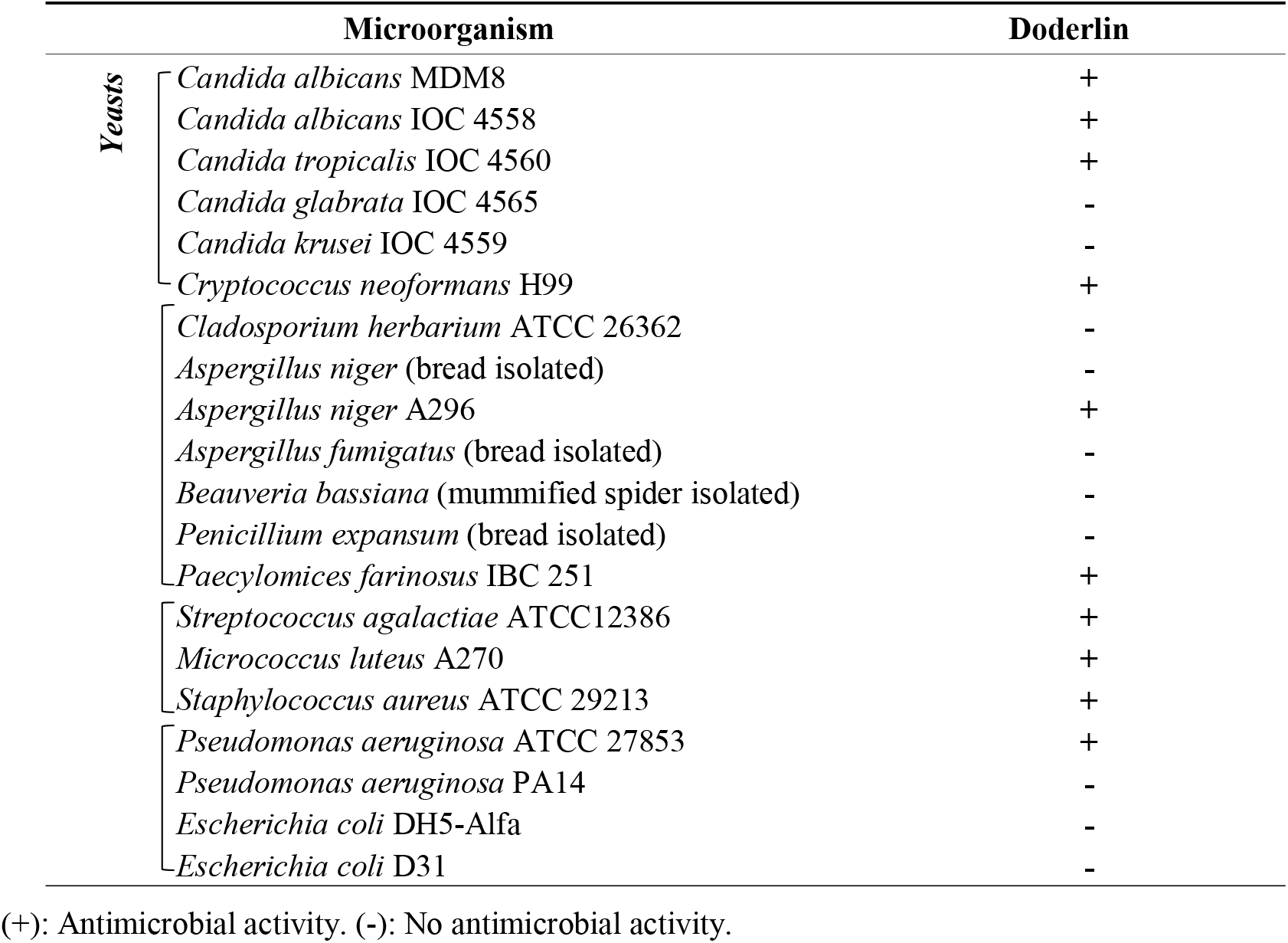
Antimicrobial activity spectrum of chemically synthesized Doderlin.

In the same experiment (2.2.1) was also possible to determine the minimum inhibitory concentration (MIC) of the Doderlin (Table 3), applying the sample in serial dilution starting from the concentration of 464 μg/mL to 29 μg/mL. Were observed that the minimum concentration to inhibit *Candida albicans IOC 4558, Candida tropicalis* IOC 4560, and *Streptococcus agalactiae* ATCC12386, were exactly corresponds to the maximum concentration applied in the test (464 μg/mL). Also, was possible to note that the most tested microorganisms had their growth inhibited at 116 μg/mL, and the lowest inhibitory concentration detected was 29 μg/mL, against the Gram-negative bacteria *Pseudomonas aeruginosa* ATCC 27853.

**Table 3.**
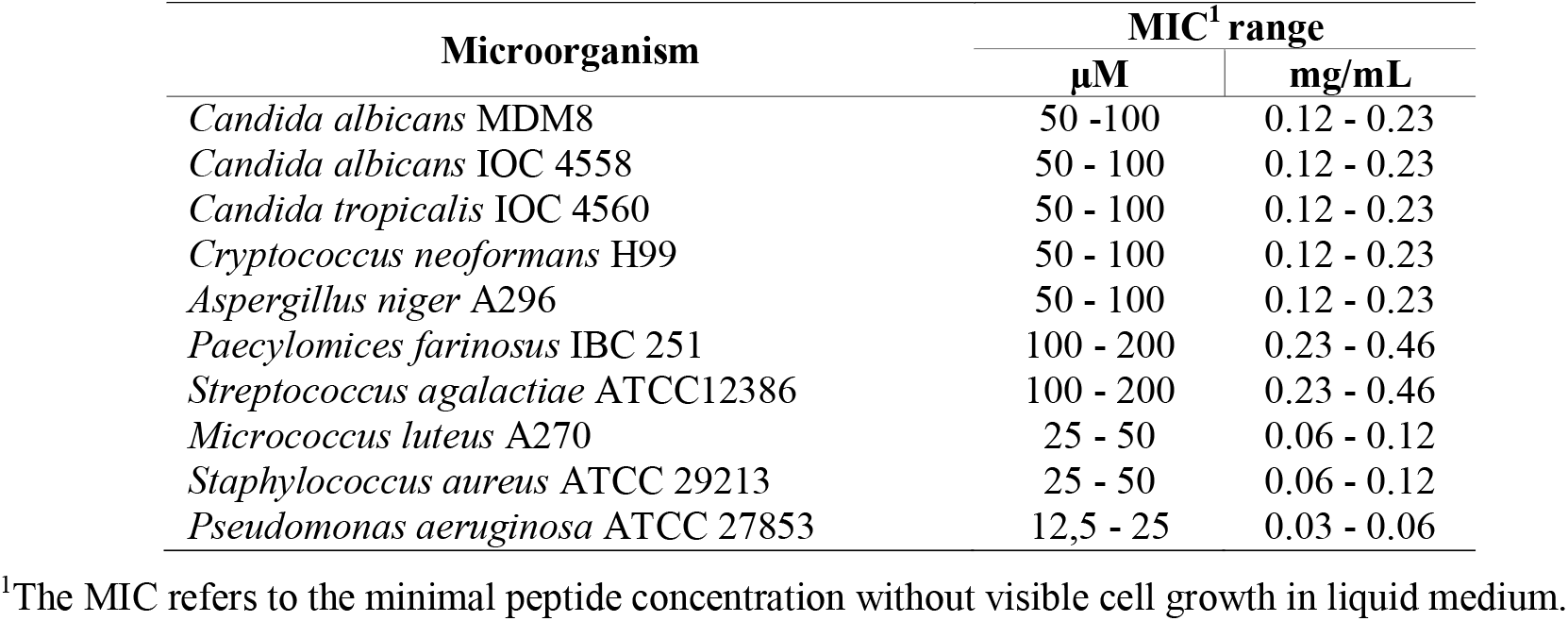
Synthetic Doderlin Minimum inhibitory concentration (MIC).

#### Hemolytic activity

Hemolytic activity was applied to determining the amount of human hemoglobin released after incubation with Doderlin. No hemolysis was observed at the evaluated concentrations, indicating that Doderlin does not have the ability to generate hemolysis in human red blood cells (Figure 7).

**Figure 7.**
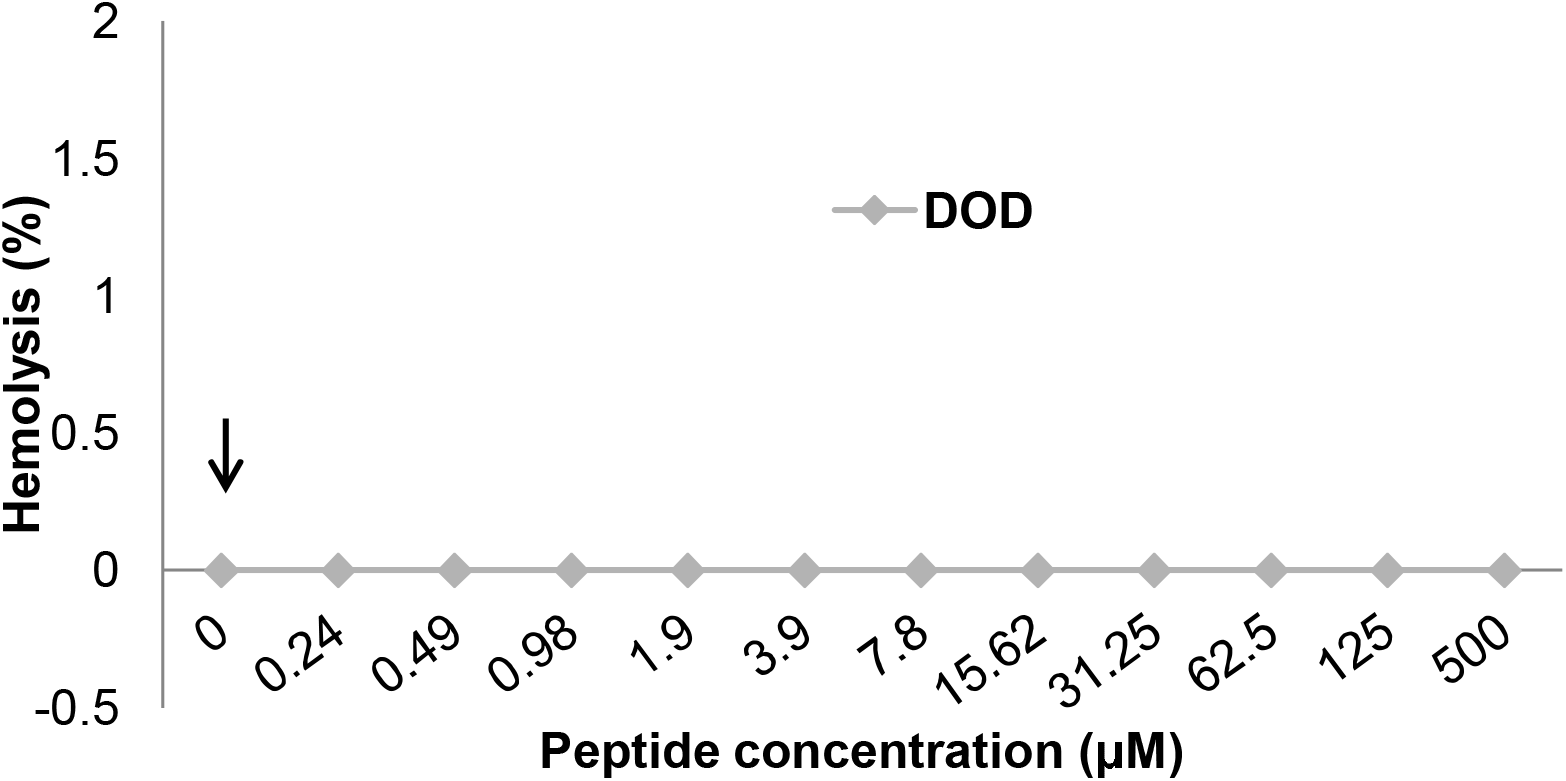
Hemolytic effects of Doderlin on human erythrocytes. Doderlin was incubated with human erythrocytes ranging from 0.24 to 500 μM for 3h at 37°C. Negative lysis control (phosphate-buffered saline - PBS) is indicated by the arrow. Positive lysis control (0.1%tritonX-100) is not indicated.

### Physicochemical properties and structural analysis

The ExPASy tool (SIB Bioinformatics Resource Portal) was used to predict the physicochemical properties of Doderlin (Table 4). This peptide was predicted to have six out of 16 hydrophobic amino acid residues, in which: two were Leu (L), two were Ala (A), and two were Phe (F) residues, with a total positive net charge (+ 1) and a high basic nature (p.I. 9.53). The Doderlin instability index was calculated, showing a value of 16.11 and suggesting that this molecule is a stable peptide (instability index < 40).

**Table 4.**
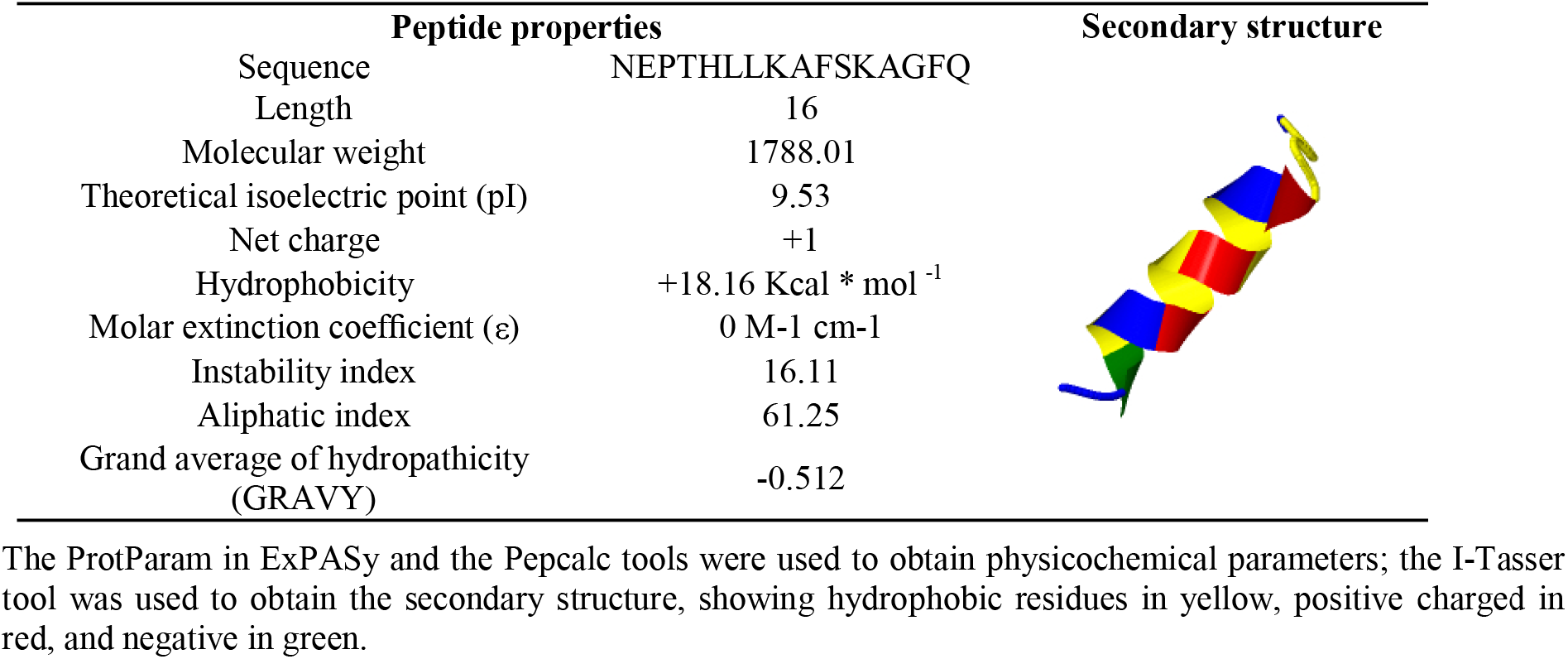
Doderlin’s theoretical physicochemical properties and secondary predicted structure.

| Peptide properties |  |
| --- | --- |
| Sequence | NEPTHLLKAFSKAGFQ |
| Length | 16 |
| Molecular weight | 1788.01 |
| Theoretical isoelectric point (pI) | 9.53 |
| Net charge | +1 |
| Hydrophobicity | +18.16 Kcal * mol<sup>-1</sup> |
| Molar extinction coefficient (ε) | 0 M-1 cm-1 |
| Instability index | 16.11 |
| Aliphatic index | 61.25 |
| Grand average of hydropathicity (GRAVY) | -0.512 |

The ProtParam in ExPASy and the Pepcalc tools were used to obtain physicochemical parameters; the I-Tasser tool was used to obtain the secondary structure, showing hydrophobic residues in yellow, positive charged in red, and negative in green.

The relative volume occupied by aliphatic side chains of a protein is termed as aliphatic index (AI). This index is important in the protein thermal stability prediction since a high AI implies in a more thermally stable protein. The AI of Doderlin is in the range of 61.25, indicating this peptide is thermally stable.

GRAVY (grand average of hydropathicity) index is related to the solubility of the proteins, indicating a hydrophobic (positive GRAVY) or hydrophilic (negative GRAVY) molecule. Doderlin presented a negative GRAVY index, suggesting it is hydrophilic and, therefore, highly soluble in water. Doderlin secondary structure was also predicted based on its primary sequence using the I-TASSER software, which showed a typical α-helix structure for this peptide (Figure 6).

## DISCUSSION

In the present study, we showed the isolation and characterization of active biomolecules from the extract of the *Lactobacillus acidophilus* cultures. Several fractions were obtained by employing the reverse phase high-performance liquid chromatography (RP-HPLC) technique, from which eight presented antimicrobial activity. These findings are in good agreement with the expected antimicrobial activity, once *Lactobacillus* sp was reported as a good source of antimicrobial peptides (AMPs) and bacteriocins (33–35).

In general, Prokaryotic AMPs have a narrow inhibitory spectrum, which is usually restricted to the microorganisms closely related to the producing bacteria (25, 36). The inhibition of growth in antimicrobial assays, as performed in this study, showed that four fractions have a more restricted antimicrobial activity, such as the fraction 3, which was effective solely against the Gram-positive *Micrococcus luteus*, and also the fractions 23, 25, and 32, which were effective only against the Gram-negative *Escherichia coli* (Table 1).

Precisely, as the antimicrobial molecules provide a competitive advantage which helps the organism to maintain its population by reducing the potential competitors, and the specificity of these peptides makes them promising antimicrobial agents with several industrial applications (31, 37). However, it is important to highlight that more tests are still needed to confirm the specificity of these molecules, since the amount obtained after the fractionation processes did not allow us to carry out that many tests necessary to clarify this point.

Besides the *L. acidophilus* antimicrobial molecules effect against Gram-positive bacteria, as expected to a lactic acid bacteria antimicrobial molecule, inhibition of Gram-negative bacteria growth was already reported (38, 39). Acidocin AA11 and Acidocin 1B are good examples of *L. acidophilus* isolated antimicrobial molecules that presents a broader spectrum of action, affecting members of the *Lactobacillus* genus and other Gram-positives bacteria, as expected to a LAB antimicrobial molecule, although not restricted to them, also affecting Gram-negative bacteria (40, 41).

Similarly, in the present study we observed the inhibition of Gram-negative bacteria, such as *E. coli* by the fractions 15, 23, 25, 29, and 32, and the inhibition of *P. aeruginosa* by the fractions 12 and 14. These microorganisms are opportunistic pathogens that causes several infections and are largely reported as multidrug resistant microorganisms (42, 43). While *E. coli* is associated to human gastrointestinal and urinary tract infections, peritonitis, bacteraemia and neonatal meningitis (44), *P. aeruginosa*, beyond respiratory infections, can also cause infections in the human urinary and gastrointestinal tracts, and bacteraemia (45), including chronic lung infections in immunocompromised individuals (46). In addition to the human infection, *P. aeruginosa* strains can also affect both livestock and pet animals, for instance, causing otitis and urinary tract infections in dogs, mastitis in dairy cows, or endometritis in horses (47). Therefore, the discovery of these new seven antimicrobial molecules has clear relevance and potential in biotechnology.

Among the eight antimicrobial fractions obtained, Doderlin (both native and synthetic version) was proven to be one of the most interesting, due to its conclusive results in chromatographic andmass spectrometry analysis, as well as its broad-spectrum inhibition, especially against *C. albicans*, which represents an important opportunistic pathogen that causes candidiasis (48). Trials using two *Lactobacillus* isolated from honey, the *L. plantarum* and the *L. curvatus*, proved the potential of these bacteria genus against several *Candida* species (49). Also, *L. acidophilus* are good producers of diverse antimicrobial substances. Some organic metabolites extracted of clinical isolates *L. acidophilus*, like lactic acid, hydrogen peroxide, and diacetyl have been tested and already demonstrated to be effective antifungal against *C. albicans* (50).

In a similar study, *Lactobacillus pentosus* TV35b isolated from vaginal secretions allowed the extraction of the Pentocin TV35b, which is a bacteriocin-like peptide with a broad inhibition spectrum with the capability to affect the *C. albicans* growth (51). Other reports also demonstrated the effectivity of antimicrobial substances against *C. albicans* using bacteriocins produced by *Streptococcus salivarius, Enterococcus faecalis*, and bacteriocins derived from protein cleavage of bovine secretion (52–54).

Despite that, this work is unique to report a native an AMP from *L. acidophilus* which is effective against *C. albicans*. This is really relevant not only because of the fact that *C. albicans* is the major responsible for *Candida* infections, but also because of the increasing number of emerging *Candida* spp resistant to azoles, such as fluconazole (FLU), the standard antifungal used for treating these infections (55, 56). In this context, Doderlin could be an important ally in combating these *Candida* infections, especially because the synthetic version of this molecule also effectively inhibited three different strains of *Candida* (Table 2).

The assays with the synthetic Doderlin were crucial because of the limited amount of native molecule obtained after the purification from the *L. acidophilus* extracts. However, these assays were consistent with the native Doderlin antimicrobial assays, once both molecules were able to inhibit the growth of the same microorganisms used at the first antimicrobial assay and allowed to expand the knowledge about the antimicrobial spectrum of Doderlin.

Another promisor antimicrobial effect observed for Doderlin was the inhibition of the *Cryptococcus neoformans* growth, which is an opportunistic yeast◻like fungal pathogen able to cause a disease that causes high morbidity and mortality in immunocompromised population, and urgently require new treatment strategies, as the cryptococcal meningitis (57), (58).

There are few reports in the literature describing AMPs with antifungal activity against *Cryptococcus*, and none of them was extracted from bacteria. However, in a following study a *de novo* designed peptide, VG16KRKP, with high potency against *C. albicans* and *C. neoformans* was developed (59). Subsequently, these authors explored the mode of action of this peptide, showing that it kills the fungal cells mainly through membrane disruption, leading to the efflux of cell content, and also using intracellular targets as a secondary mode of action (60). The mode of action of Doderlin still need further investigation, but giving its physicochemical characteristics, as described here, it is possible to hypothesize that this peptide may also act through these two modes of action as well.

Regarding to the inhibition of bacterial growth, the results obtained here for Doderlin were similar to those previously reported for a bacteriocin from the extracts of the *L. acidophilus* ks400, which was effective against *Streptococcus agalactiae* and *Pseudomonas aeruginosa* (61), as noticed for Doderlin herein. Moreover, the antimicrobial activity of Doderlin against *Streptococcus agalactiae* is very relevant, considering that this is one of the most relevant opportunistic pathogens related to vaginal infections and also the leading cause of neonatal disease worldwide (62).

On one hand, synthetic Doderlin assays allowed to expand the knowledge about its antimicrobial spectrum, and on the other, relative high concentration of the peptide was required to affect the growth of tested microorganisms.

In general, the majority of previous works described very low MIC values for antimicrobial molecules (63, 64), in which bacteriocins isolated from *Lactobacillus* displayed MIC ranges from 0.8 μg/mL to 23 ng/mL against diverse microorganisms. Nevertheless, comparing the MIC values of Doderlin, against similar microorganisms, to some other *Lactobacillus* broad-spectrum antimicrobial substances, it seems that the MIC values do not differ significantly. An example is the comparation of the MIC values necessary to the *S. aureus* and *M. luteus* growth inhibition. The NX2-6 bacteriocin inhibited the growth of mentioned bacteria at a concentration of 80 and 120 μg/mL, respectively (65), while Doderlin MIC were 116 μg/mL for both bacteria. Moreover, a bacteriocin like-substance from *L. plantarum*, the LBP102, also inhibited *S. aureus* at the concentration of 70 μg/mL (66), which is not considerably lower than the Doderlin minimum concentration.

Surprisingly, the MIC of Doderlin against *C. albicans* was one of the highest observed (above 200 μg/mL), whereas the most notable Doderlin MIC value was against *P. aeruginosa* at 29 μg/mL, which was lower than the 70 μg/mL observed in the study with the antimicrobial substance LBP102 (66). Indeed, this MIC value for Doderlin is still low even if compared to some AMPs, like the OVTP12, a peptide derived from egg tested against the same *P. aeruginosa* strain used in the present study, but effective in a higher concentration (128 μg/mL) than Doderlin (67).

Some physicochemical parameters of Doderlin were also explored. Indeed, physicochemical properties and the sequence of peptides can directly affect their functions, bioavailability and bioaccessibility, once that they are released from the precursor protein where they are encrypted (68). The MW of the peptide NEPTHLLKAFSKAGFQ is 1788.01 Da, and the net charge was predicted to be positive, a known evidence of membrane electrostatic interaction, that facilitates the initial binding of the positively charged peptides to the negatively charged bacterial membrane (69). Similarly, peptide hydrophobicity will also influence the uptake and bioactivity of peptides (70).

The instability index (II) predicted classifies Doderlin as a stable peptide (71). PepDraw server predicted the hydrophobicity of the peptide Doderlin as +18.16 kcal mol^−1^ with the Wimley-White scale. This peptide has a good solubility in water, and the grand average of hydropathicity (GRAVY) of this peptide is −0.512, which indicates that this peptide is hydrophilic; these properties were also previously observed for the AVPYPQR novel casein anticoagulant peptide (https://www.ncbi.nlm.nih.gov/pubmed/30994118). Also, Doderlin predicted secondary structure shows an α-helix that is commonly reported as a characteristic of AMPs, that usually acts on the bacteria membrane, in the same way as the human cathelicidin LL-37 (72) and magainin-2 (73).

## CONCLUSIONS

The present work allowed the isolation and characterization of novel antimicrobial peptide (AMP) which was extracted from *Lactobacillus acidophilus* and showed a broad antimicrobial spectrum (against Gram-positive and negative bacteria, and yeast). The most promising molecule was purified and characterized, and named Doderlin. Doderlin is more effective against Gram-positive bacteria and yeasts with a minimum inhibitory concentration (MIC) of 116 μg/mL for the majority of tested microorganisms. Even though, *P. aeruginosa* was the most sensible bacteria to Doderlin, with a MIC value of only 29 μg/mL. In addition, the bioinformatic analysis predict Doderlin as a stable and soluble in water peptide, with α-helix secondary structure, suggesting it may interact with lipid membranes. These findings make this molecule very interesting from a biotechnological point of view and a potential antimicrobial candidate to be applied in food and pharmaceutical industry. Complementary studies are required to continue the characterization of the other seven antimicrobial fractions detected, and to compare the differences in efficacy and potency between native and synthetic Doderlin versions. Also, the mode of action of the peptide needs to be further investigated.

## MATERIALS AND METHODS

### Bacterial strains and growth conditions

The *L. acidophilus* strains were collected from the vagina of patients at the São Paulo Hospital, located at Federal University of São Paulo (UNIFESP). The strains isolation and identification were also performed by the Hospital’s coworkers. Samples were transported on ice to the Special Laboratory for Applied Toxinology (LETA) at Butantan Institute (São Paulo, Brazil), and stored at −80 °C in MRS broth containing 20% (vol/vol) glycerol. *The clinical isolated Lactobacillus acidophilus* were cultivated in 10L of the growth media recommended, and kept under agitation at 37°C for 24 h, without aeration until mid-logarithmic phase of growth (O.D. _600_ = 0.8).

Bacterial and fungal strains used in the antimicrobial assays were: *Candida albicans* MDM8, *Candida albicans* IOC 4558, *Candida tropicalis* IOC 4560, *Candida glabrata* IOC 4565, *Candida krusei* IOC 4559, *Cryptococcus neoformans* H99, *Cladosporium herbarium* ATCC 26362, *Aspergillus niger, Aspergillus fumigatus*, and *Penicillium expansum* (bread isolated*), Aspergillus niger* A296, *Paecylomices farinosus* IBC 251, *Streptococcus agalactiae* ATCC12386, *Micrococcus luteus* A270, *Staphylococcus aureus, Pseudomonas aeruginosa* ATCC 27853, *Pseudomonas aeruginosa* PA14, *Escherichia coli* DH5-alfa, *Escherichia coli* D31, *Escherichia coli SBS363* and mummified spider isolated *Beauveria bassiana*, belonging to the collection of microorganisms of the Special Laboratory for Applied Toxinology (LETA) of the Butantan Institute (São Paulo, Brazil).

### Production of culture supernatants

After incubation, the whole broth was centrifuged at 2,110 × g for 15 min at 4 °C, for cells pelletization. The cells pellets were resuspended in a buffer containing 200 mL of 2M acetic acid, mixed in a magnetic stirrer for 30 min in the presence of ice. The suspension was lysed by ultracentrifugation for 16,000 ×g for 30 min at 4 °C to obtain the supernatant.

#### Solid Phase Extraction

The supernatant was partially purified by Sep-Pak C18 cartridges (Waters Associates, Milford, MA, USA), using three successive acetonitrile (ACN) concentrations in water (5, 40, and 80%). The eluted samples were lyophilized and reconstituted in 2 ml trifluoroacetic acid (0.05% TFA). Then, it was centrifugated at 14,000 ×g for 2 min to remove the insoluble material before submitting the material to liquid chromatography.

#### Peptide fractionation

The supernatant was fractionated by a reverse-phase high-performance liquid chromatography (RP-HPLC) at room temperature on a Shimadzu LC-10 HPLC system using a semi-preparative C18 Jupiter column (10 mm; 300A; 10 × 250 mm) (Phenomenex International, Torrance, CA, USA) equilibrated at room temperature with 0.05% TFA in ultrapure water. The elution was done under a linear ACN gradient and the flow rate was 1.5 mL/min during 60 min (0-20% for fractions eluted in 5%, 2 - 60% for fractions eluted in 40%, and 20-80% for fractions eluted in 80%). Each fraction was individually and manually collected, under Ultraviolet (74) absorbance monitored at 225 nm. The fractions were lyophilized, reconstituted in 500 μL ultrapure water, and used in antimicrobial activity assays.

The two fractions with antibacterial activity (12 and 29) were submitted to a second RP-HPLC step (1 mL/min flow rate during 60 min) using an analytical C18 Jupiter column (10 mm; 300A; 4.6 × 250 mm) (Phenomenex International, Torrance, CA, USA). The ACN concentration ranged from 0% to 10% (fraction 12) and from 19 to 29% (fraction 29). The antibacterial activity of each fraction was then tested again.

Size-exclusion chromatography (SEC) was employed to the Doderlin isolation in an ÄKTA purifier 10 (GE Healthcare, Chicago, Illinois, EUA) system, using a Superdexpeptide HR 10/30 (7.5 × 300 mm) (GE Healthcare) column. The sample was reconstituted in ammonium acetate (50 mM) and centrifugated at 16,000 ×g, for 5 min. Then, the supernatant was submitted to the system under a 1 mL/min flow rate during 36 min and monitoring the absorbance at 280 nm. Eluted fractions were every minute automatically collected and immediately refrigerated, concentrated in a vacuum centrifuge SpeedVac Savant (Thermo Fisher Scientific, Waltham, MA, USA) and stored in the *freezer* at −20 °C until its usage.

#### Antimicrobial Assays

The antimicrobial activities of the fractions were evaluated by a liquid growth inhibition assay (75, 76). The assay was performed using a serial dilution in 96-wells sterile plates, where each well was filled with 20 μL of the fraction and 80 μL of bacterial dilution, providing a final volume of 100 μL (77). Bacteria were cultured in poor nutrient broth (PB) (1.0 g peptone in 100 ml of water containing 86 mM NaCl at pH 7.4; 217 mOsm), and the fungi and yeast were cultured in a poor potato dextrose broth (1/2-strength PDB) (1.2 g potato dextrose in 100 mL of water at pH 5.0; 79 mOsm). Exponential growth phase cultures were diluted to 5 × 10^4^ CFU/mL final concentration. Sterile water was used as growth control, and streptomycin or tetracycline was used as growth inhibition control. Plates were incubated for 18 h (bacteria) or 24 h (fungi/yeast) at 30 °C. Growth inhibition was determined by measuring absorbance at 595 nm using the equipment Victor3 (Perkin Elmer Inc.). Fractions were tested in triplicate.

The Minimum Inhibitory Concentration (MIC) was established according to the previously described (Section 4.3 and 4.4), using the fraction 12 (Doderlin) in serial dilutions starting from the initial concentration of 200 μM (432 μg/mL) against Gram-negative bacterial strains, Gram-positive bacterial s, fungal, and yeast strains. The MIC was defined as the lowest concentration of sample that caused 100% growth inhibitions, measured by Victor3 spectrophotometer (Perkin Elmer Inc.) at 595 nm.

#### Hemolytic assay

Hemolytic activity of Doderlin was determined using fresh human red blood cells (hRBCs) donated by a healthy adult, washed three times with phosphate-buffered saline (PBS) (35 mM phosphate buffer, 0.15 M NaCl, pH 7.4) by centrifugations at 700 ×g for five minutes to a final 3% (vol/vol) concentration. Doderlin solutions were added to 50 μL hRBC suspension to a final 100 μl and incubated in a U-shaped bottom plate for 3 h at 37 °C. The sample concentration applied started from 500 μM (1160 μg/mL) to 0.24 μM (0.557 μg/mL), in serial dilution.

Release of hemoglobin was determined by measuring the supernatant absorbance (Abs) at 405 nm and calculated as a percentage of 100% lysis control (0.1% Triton X-100) (Sigma-Aldrich, St. Louis, MO, USA), following the equation: % hemolysis = (Abs sample - negative Abs)/(positive Abs - negative Abs). PBS was used as a negative control (78, 79). Assays were done in triplicate. The Ethics Committee was the University of Sao Paulo School of Medicine (USP), N° 21884919.5.0000.0065.

#### Mass spectrometry analysis and Doderlin identification

Active antibacterial fractions (reconstituted in formic acid 0.1%), were analysed on an LTQ-Orbitrap Velos (Thermo Scientific) mass spectrometry coupled to an Easy-nLC II liquid nano-chromatography (Thermo Scientific). Ten μL of each sample were automatically injected on a C18 pre-column (100 mm I.D. × 50 mm; Jupiter 10 mm, Phenomenex Inc., Torrance, California, USA) coupled to a C18 analytical column (75 mm I.D. × 100 mm; ACQUA 5 mm, Phenomenex Inc.), at a flow rate of 200 nL/min under a linear gradient (from 0 to 95% mobile phase B) (0.1% formic acid in 100% acetonitrile) for 15 min. The electrospray source was operated at 2 kV and 200 °C in positive ion mode. Mass spectra were acquired by Fourier Transform Mass Spectrometry (FTMS) analyser in full scan mode (MS), in the range of 200 to 2,000 m/z with a resolution of 60,000 at 400 m/z. The 10 most intense peaks were automatically selected via data dependent acquisition for the subsequent acquisition of the spectra of the product ions (MS/MS). The minimum threshold for selecting an ion for a fragmentation event (MS/MS) was set to 5,000 counts per second (cps), and the dynamic exclusion time was set to 30 s. Normalized collision energy was set to 35%.

### Synthetic Doderlin peptide

Doderlin (amino acid sequence: NEPTHLLKAFSKAGFQ) was synthesized by solid phase synthesis at China Peptides Co. Ltd (Shanghai, China), with purity of 70%, not amidated.

### Doderlin bioinformatic analysis

The MS/MS peak list files were submitted to an in-house version of the MASCOT server (Matrix Science, USA) and screened against the NCBI and SwissProt databases. For mass analysis by the deconvolution of the ions was used the MagTran® Program, version 1.02 (80).

The primary structure of the Doderlin was drawn using the PepDraw (http://pepdraw.com/) peptide calculator.

Doderlin physicochemical characteristics were evaluated using online peptide calculators. The theoretical molecular weight, isoelectric point, the peptide charge at pH 7 and extinction coefficient of the peptide were estimated with the online Pepcalc software (http://pepcalc.com/), the Instability and Aliphatic index was estimated with the ProtParam software (https://web.expasy.org/protparam/), and the I-Tasser tool (https://zhanglab.ccmb.med.umich.edu/I-TASSER/) was used to obtain the Doderlin predicted secondary structure.

## ACKNOWLEDGMENTS

We thank all the team of the Protein Chemistry Laboratory at the Special Toxinology Laboratory (LETA-Butantan Institute, Brazil) for the constant emotional support and encouragement, in special to Msc. Soraia Maria do Nascimento, Msc. Thiago de Jesus Oliveira, Msc Sandra Regina dos Santos, Dr. Débora Álvares Leite Figueiredo and Dr. Katie Cristina Takeuti Riciluca. Also, we thank the technicians Rosa Maria Carmo, Ismael Feitosa Lima, Ivan Novaski Avino and Lídia Alves da Silva.

## Author Contributions

Conceptualization, B.S.S., M.A.F.H. and P.I.S.J.; methodology B.S.S, E.SY, M.A.F.H. and P.I.S.J.; software, A.D.R., B.S.S and P.I.S.J.; validation, B.S.S, E.SY, M.A.F.H. and P.I.S.J.; formal analysis, A.D.R., B.S.S and P.I.S.J.; investigation, B.S.S, E.SY, M.A.F.H. and P.I.S.J.; resources, P.I.S.J.; data curation, A.D.R., B.S.S and P.I.S.J.; writing—original draft preparation, A.D.R. and B.S.S; writing, review and editing, A.D.R., B.S.S, M.A.F.H. and P.I.S.J.; supervision, P.I.S.J.; project ad-ministration, P.I.S.J.; funding acquisition, P.I.S.J. All authors have read and agreed to the pub-lished version of the manuscript.

